# Isolation and semi quantitative PCR of Na^+^/H^+^ antiporter (SOS1& NHX) genes under salinity stress in *Kochia scoparia*

**DOI:** 10.1101/145953

**Authors:** Leila fahmideh, Ziba Fooladvand

## Abstract

*Kochia scoparia* is a dicotyledonous annual herb and belongs to the Amaranthaceae family. Genetic diversity and resistance to drought stress of this plant has made it widely scattered in different regions which contains highly genetic diversity and great potential as fodder and can grow on salty, drought affected areas. Since the soil salinity has become widely spread. environmental concern has sparked so many debates. An important limiting factor in agricultural production worldwide is the sensitivity of most of the crop to salinity caused by high concentration of salts soil. Plants use three different strategies to prevent and adapt to high Na^+^ concentrations. Antiporters are important category of genes that play a pivotal role in ion homeostasis in plants. Na^+^/H^+^ antiporters (NHX1 and SOS1) are located in tonoplasts and reduce cytosolic Na^+^ concentration by pumping in the vacuole whereas SOS1 is localized at the plasma membrane and extrudes Na^+^ in apoplasts. Coding sequence of plasma membrane Na ^+^/H ^+^ antiporter (SOS1) and vacuole membrane Na /^+^H ^+^ antiporter (NHX) in *Kochia scoparia* were isolated using conserved sequences of SOS1 and NHX. Also, expression profile under salinity stress was studied in this study. The amino acid sequences (aa) of the isolated region of K.SSOS1 and K.SNHX showed the maximum identity up to 84% and 90% to its orthologue in salicornia brachiata and suadea maritima, respectively. The results of semi-quantitative RT-PCR revealed that salinization has affected positively on SOS1 transcription level. The expression of K.SSOS1 and K.SNHX in leaves and roots of *Kochia scoparia* were progressively increased under all salinity levels compared to control. The results suggest that K.SSOS1 and K.SNHX play an essential role in salt tolerance of *K.scoparia* and they can be useful to improve salt tolerance in other crops.

## Introduction

Most of studies have revealed that the greatest lost in various crop production is due to abiotic stresses, such as, salinity, water deficit, low temperature and heavy metals adversely affect the growth and several physiological processes such as leaf cell growth and biomass production of plants. An important limiting factor in agricultural production worldwide is the sensitivity of most of the crop to salinity caused by high concentration of salts soil. Processes such as seed germination, seedling growth and vigor, vegetative growth, flowering and fruit set are adversely affected by high salt concentration, ultimately causing yield lost. Salinity stress can reduce the productivity of glycophytes, which are the majority of agricultural products. High salt concentrations cause hyper osmotic stress and ion imbalance in plants which often as a secondary effect leads to oxidative damage in cellular components (Qadir, Tubeileh et al. 2008). Plants adapt to environmental stresses via responses, including the activation of molecular networks that regulate stress perception, signal transduction and the expression of both stress related genes and metabolites (Huang, Ma et al. 2012). Plants have stress specific adaptive responses as well as responses which protect the plants from more than one environmental stress (Huang, Ma et al. 2012). Plants employ three different strategies to prevent and adapt to high Na^+^ concentrations: 1) active Na^+^ efflux, 2) Na^+^ compartmentalization in vacuoles, and 3) Na^+^ influx prevention (Niu, Bressan et al. 1995, Rajendran, Tester et al. 2009). Antiporters are important groups of genes that have a key role in ion homeostasis in plants. Na^+^/H^+^ antiporters (NHX1 and SOS1) maintain the appropriate concentration of ions in the cytosol, thereby minimizing cytotoxicity. NHX1 are located in tonoplasts and reduce cytosolic Na^+^ concentration by pumping it in the vacuole (Gaxiola, Rao et al. 1999), whereas SOS1 is localized at the plasma membrane and extrudes Na^+^ in apoplasts (Shi, Quintero et al. 2002). Both of these antiporters are driven by a motive proton force generated by the H^+^-ATPase (Blumwald, Aharon et al. 2000). The SOS signaling pathway consists of three major proteins including: SOS1, SOS2, and SOS3. SOS1, which encodes a plasma membrane Na ^+^/H ^+^ antiporter, is essential in regulating Na^+^ efflux at the cellular level. It also facilitates long distance transportation of Na^+^ from root to shoot. Over expression of this protein leads to salt tolerance in plants (Shi, Ishitani et al. 2000). Activation of SOS1 by direct phosphorylation of the self-regulation scope is possible by serine/threonine protein kinas or SOS2 that requires calcium binding protein or SOS3 (Quintero, Martinez-Atienza et al. 2011). C-terminal end of the protein causes the Na^+^ to move. At the C-terminal end, SOS1, the 764 849 region is cyclic nucleotide-binding site and in the 998 -1146 region a self-regulator domain exists. In the respite state the self-regulator domain interacts with upstream sequence bearing the cyclic nucleotide-binding site (Quintero, Martinez-Atienza et al. 2011). In fact, the self-regulator domain is a target location for phosphorylation by SOS2. After SOS1 phosphorylation, the self-regulator domain leaves upstream location and attaches at this location of cyclic nucleotide and transferring protein activity begins (Quintero, Martinez-Atienza et al. 2011). According to the above mentioned information, domain connected to the cyclic nucleotide can be used as one of the most important locations to regulate SOS1 activity, eventually its effect on salinity tolerance.

*K.scoparia*, a dicotyledonous erect annual herb belongs to Amaranthaceae family with high genetic diversity and great foliage potential (Mullinex 1998), reported that *K scoparia* on of its Iranian variety is highly tolerant to salt and could be considered as a foliage species in cold regions of the world. Rapid vegetative growth under high salinity and temperature and drought and stress makes it a very valuable candidate as a non-conventional foliage crop for arid temperate regions (Kafi, Asadi et al. 2010). *K scoparia* has been widely used in Chinese and Korean traditional medicine as a treatment for skin diseases, diabetes, mellitus, rheumatoid arthritis, liver disorders, and jaundice (Choi, Lee et al. 2002, Kim, Lee et al. 2005). Kochia seeds contain an ovi position pheromone that can be added as an attractant for mosquito pesticides (Whitney, Sayanova et al. 2004, Friesen, Beckie et al. 2009). It has been reported that seeds of Kochia also contain other chemicals that could be beneficial for human, such as compounds used in ulcers, rheumatoid arthritis, treatmeant and some pathogenic bacteria (Friesen, Beckie et al. 2009) (Friesen et al., 2009; Goyal and Gupta, 1988; Borrelli and Izzo, 2000). The aim of this study was to investigate the presence of SOS1 and NHX1 genes and trace it using by induced salt stress in *Kochia scoparia*, Futures of these genes in protein structure characterized with in silico tools. Furthermore profiling gene expression for two gene charecterized. *K.scoparia* is an attractive plant model for study the mechanism of salt tolerance. This work to gain insights into the role played by this transporter in *K.scoparia* halophyte.

## Material and methods

### Genetic samples

*K.scoparia* was collected from Sabzevar in Khorasan Razavi Agricultural Research Center (Iran) and planted in Biotechnology research Center University of Zabol. The fresh leaves were applied to isolate RNA after salinity stress(plants were irrigated by 100mM, 200mM, 300mM and 400mM sodium chloride solutions).

### Primers design

17 SOS1 and 22 NHX coding sequences data which are available at NCBI data base(ncbi.nlm.nih.gov) have been showed in Table(1),were aligned by ClustalW method provided in DNASTAR Laser gene software (EditSeq, Meg-Align, Version 5.00), GENEDOC (Multiple Sequence Alignment Editor & Shading Utility Version 2.5.000). all specific primers designed based on the most conservative parts of the alignments. specific forward and reverse primers were designed (Table 2).

**Table 1:**
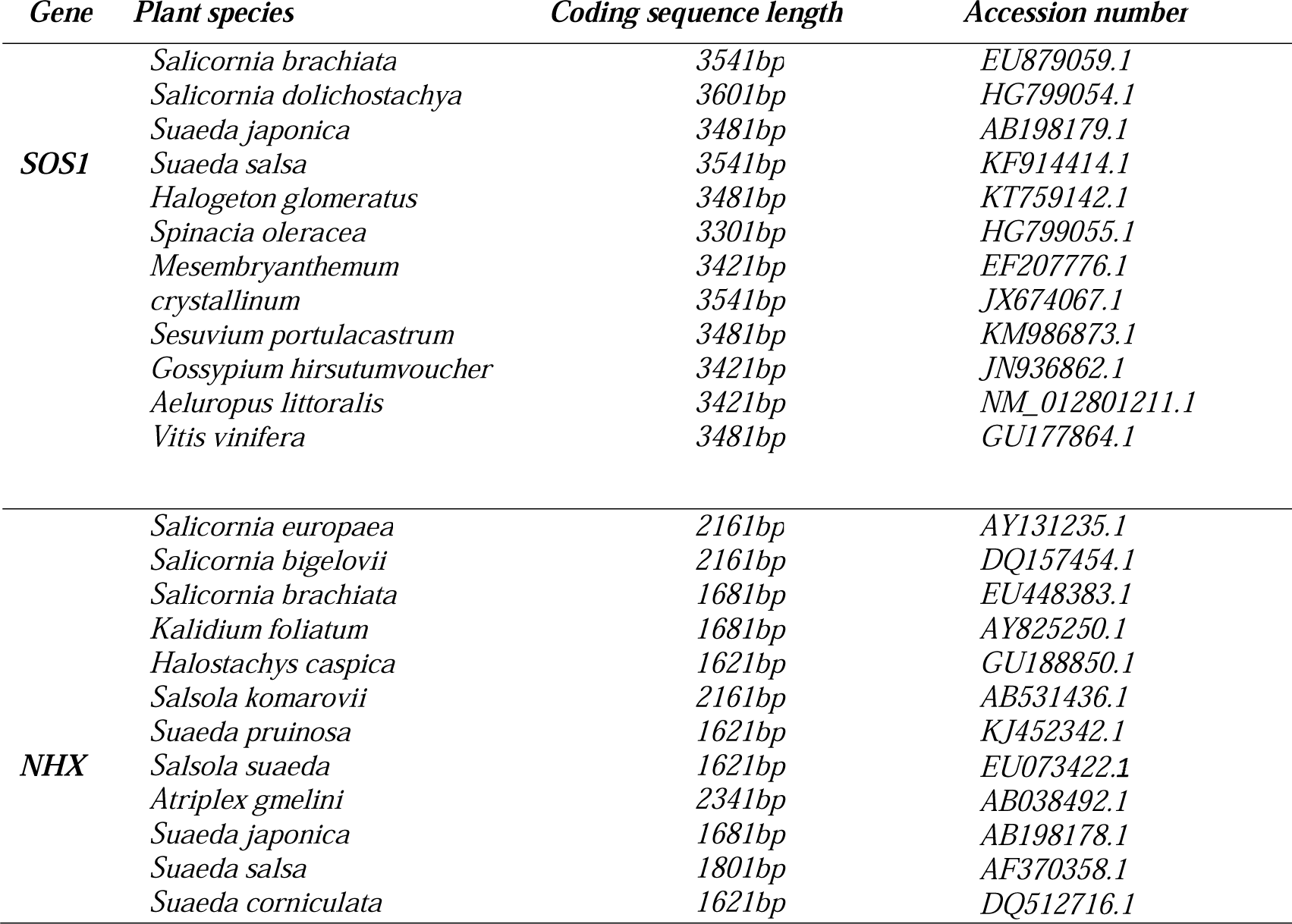
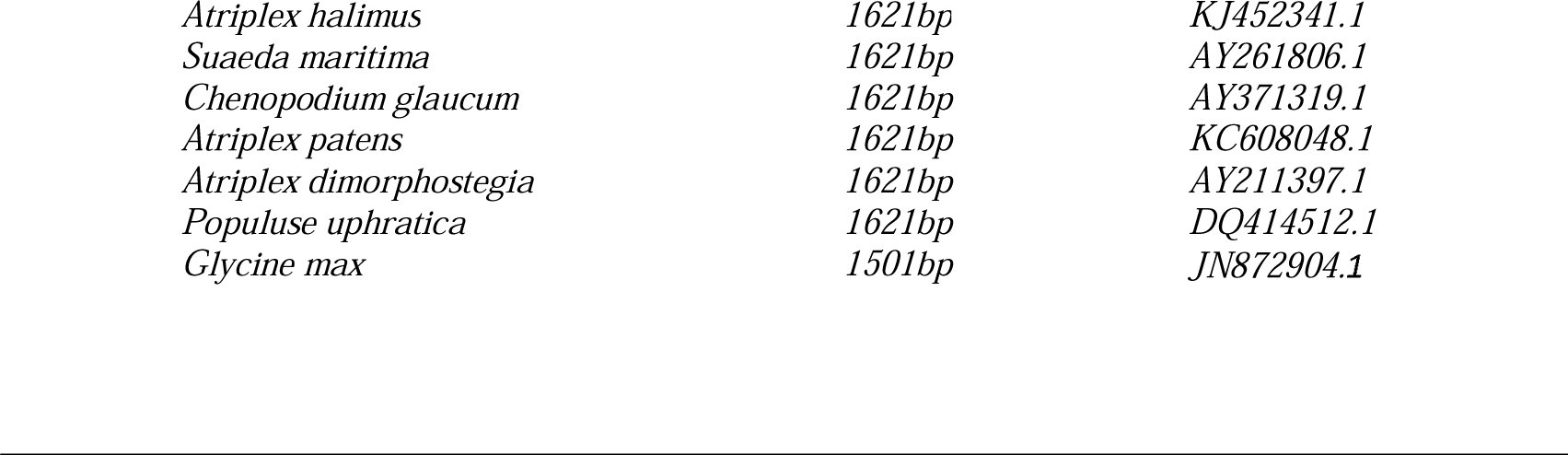
Plant species and accession numbers of gene sequences used for primer desi-gn alignment.

| *Gene* | *Plant species* | *Coding sequence length* | *Accession number* |
| --- | --- | --- | --- |
| **SOS1** | *Salicornia brachiata* | 3541bp | EU879059.1 |
|  | *Salicornia dolichostachya* | 3601bp | HG799054.1 |
|  | *Suaeda japonica* | 3481bp | AB198179.1 |
|  | *Suaeda salsa* | 3541bp | KF914414.1 |
|  | *Halogeton glomeratus* | 3481bp | KT759142.1 |
|  | *Spinacia oleracea* | 3301bp | HG799055.1 |
|  | *Mesembryanthemum* | 3421bp | EF207776.1 |
|  | *crystallinum* | 3541bp | JX674067.1 |
|  | *Sesuvium portulacastrum* | 3481bp | KM986873.1 |
|  | *Gossypium hirsutumvoucher* | 3421bp | JN936862.1 |
|  | *Aeluropus littoralis* | 3421bp | NM_012801211.1 |
|  | *Vitis vinifera* | 3481bp | GU177864.1 |
| **NHX** | *Salicornia europaea* | 2161bp | AY131235.1 |
|  | *Salicornia bigelovii* | 2161bp | DQ157454.1 |
|  | *Salicornia brachiata* | 1681bp | EU448383.1 |
|  | *Kalidium foliatum* | 1681bp | AY825250.1 |
|  | *Halostachys caspica* | 1621bp | GU188850.1 |
|  | *Salsola komarovii* | 2161bp | AB531436.1 |
|  | *Suaeda pruinosa* | 1621bp | KJ452342.1 |
|  | *Salsola suaeda* | 1621bp | EU073422.1 |
|  | *Atriplex gmelini* | 2341bp | AB038492.1 |
|  | *Suaeda japonica* | 1681bp | AB198178.1 |
|  | *Suaeda salsa* | 1801bp | AF370358.1 |
|  | *Suaeda corniculata* | 1621bp | DQ512716.1 |

|  |  |  |
| --- | --- | --- |
| *Atriplex halimus* | 1621bp | KJ452341.1 |
| *Suaeda maritima* | 1621bp | AY261806.1 |
| *Chenopodium glaucum* | 1621bp | AY371319.1 |
| *Atriplex patens* | 1621bp | KC608048.1 |
| *Atriplex dimorphostegia* | 1621bp | AY211397.1 |
| *Populus euphratica* | 1621bp | DQ414512.1 |
| *Glycine max* | 1501bp | JN872904.1 |

**Table 2.**
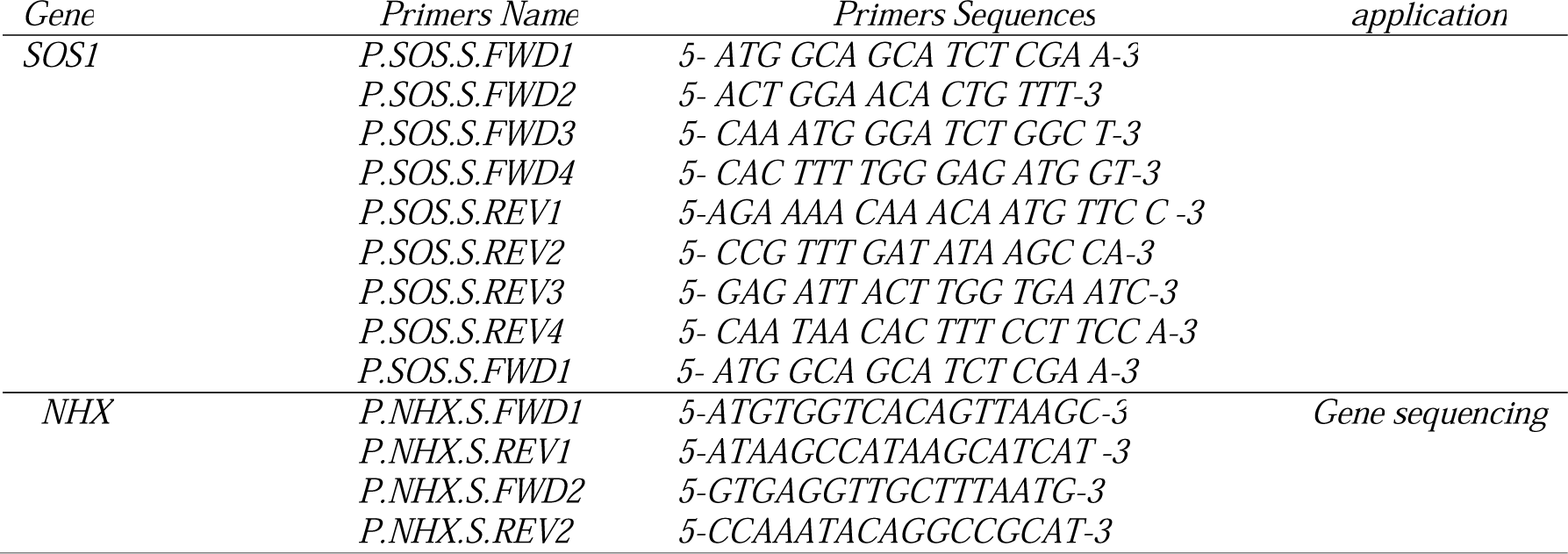
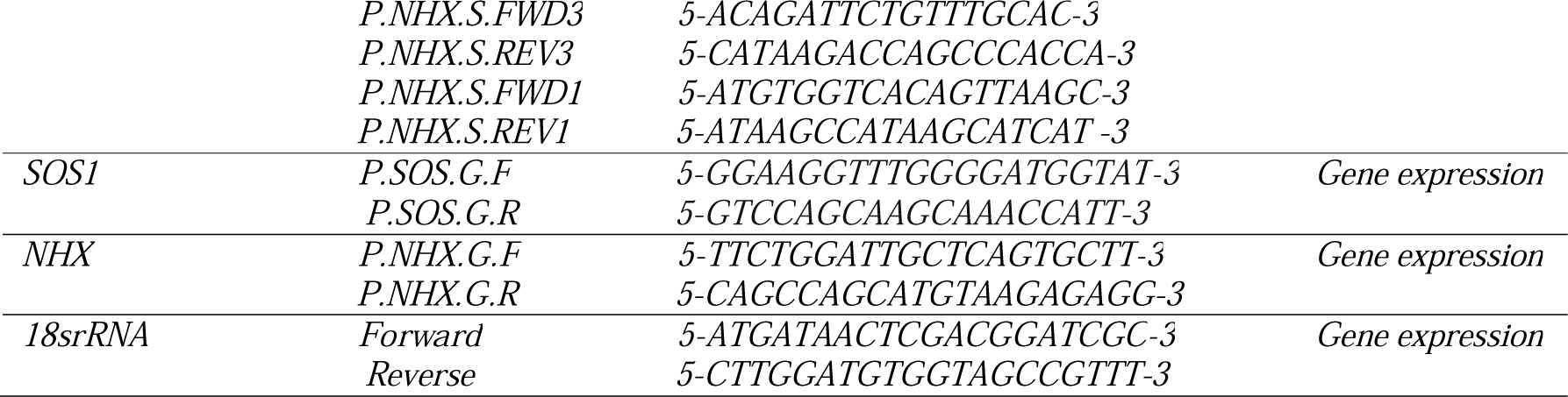
Primers sequences and names were developed for isolation and gene expression of SOS1, NHX in *K.scoparia.*

### RNA isolation and cDNA amplification in *K.scoparia*

Total RNA of samples were isolated by Total RNA isolation kit (DENA Zist Asia). The cDNA(s) were synthesized using Hyper script reverse transcriptase (Gene All) and oligod (T) 18mer, P.SOS.S-REV1, P.SOS.S-REV2, P.SOS.S-REV3, P.SOS.S-REV4, P.NHX.S.REV1, P.NHX.S.REV2 and P.NHX.S.REV3primers (Table1) and amplified with a combination of primers (Table 2). The amplifications were obtained in 30 cycles at defined annealing temperature for each pair of primers using TaqDNA polymerase (AMPLIQON). The process finished after a final extension for 5-15 min at 72°C (Fig.1).

**Figure 1:**
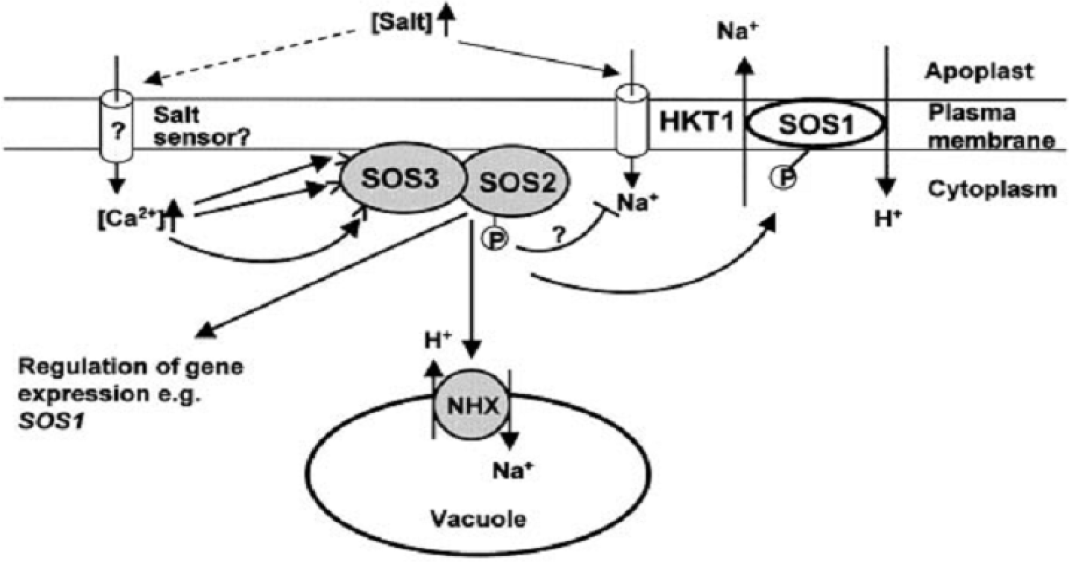
Regulation of ion homeostasis by ion Na^+^/H^+^ pumps antiporters (SOS1), vacuolar Na^+^/H^+^ exchanger (NHX) that salt sensors present at the plasma and vacuolar membranes (Chinnusamy, Schumaker et al. 2004).

### Sequencing of SOS1 and NHX in *K.scoparia*

PCR products were extracted and purified from 0.8% agarose gel using GEL recovery DNA kit (DENA Zist Asia). PCR reactions were sequenced utilizing Euro fins MWG Operon company service. Sequence analysis including deletion of error in sequences, assembly of fragments, alignment with other plant species gene sequences, was done using DNA STAR Laser gene software (EditSeq, SeqManII Meg-Align, MapDraw; Version5.00), GENEDOC (Multiple Sequence Alignment Editor & Shading Utility Version 2.5.000 and NCBI BLAST (Altschul, Gish et al 1990). The amino acid sequences were aligned with CLUSTALW software. SOS1 and NHX nucleotides and amino acid sequences aligned and analyzed with SOS1 and NHX from other plants (Fig.3). Assembled sequences for gene submitted to DDBJ data base addressed: ddbj.nig.ac.jp and assigned accession number (LC218450, LC218451 for SOS1 and NHX gene respectively).

### RT-PCR for analyzing of SOS1 and NHX gene expression in *K.scoparia* under salt stress

Eight week old seedlings Plants were treated in Salinity stress. Afterward plants were irrigated by 150 mM, 300mM sodium chloride solutions. Then, sampling was done during 12, 48, 72 hours after treatment. RNAs were extracted using Total RNA isolation kit (DENA Zist Asia, Iran) from the treated seedlings according to manufacture^S^ instructions. After Dnase1 treatment of RNA samples, 2µg of RNAs, using Gene All first strand cDNA Synthesis Kit, was reverse transcribed to cDNAs, that were used as templates for semi quantitative RT-PCR. The cDNA amounts were first normalized by 18s rRNA PCR product intensity. PCR process was performed using the following procedure: 95 °C for 5 min followed by 35 cycles of 95°C for 30 sec, annealing temperature for 45sec, and 72°C for 1 min, and finally 15 min at 72°C for final extension.

### Gel analysis for gene expression of SOS1 and NHX in *K.scoparia* under salt stress

Images of the RT-PCR ethidium bromide-stained agarose gels were taken with a Vilber documentation system (E-BOX CX5) and Band intensity was expressed as relative absorbance units. The ratio between the sample Total RNA and 18srRNA was determined and calculated to normalize for initial variations in sample concentration and as a control for reaction efficiency. Mean and standard deviation of all experiments were calculated after normalization to 18srRNA.

### Molecular docking to predict SOS1 protein of *K.scoparia* and regulation in the salt stress

Molecular docking of the desired fragments isolated from Kochia using SWISSDOCK was performed as followed percedure: First, tertiary structure of sequence fragments was predicted in the by SWISS-MODEL is a fully automated protein structure homology - modelling server (Arnold, Bordoli et al. 2006, Guex, Peitsch et al. 2009, Kiefer, Arnold et al. 2009, Biasini, Bienert et al. 2014). Then, target ligand, cAMP from Zinc dock using online service of Swiss Dock available on the Expasy site molecular docking. The best state for interaction was reported using UCSF-Chimera method (Bordoli, Kiefer et al. 2009).

## Results and Discussions

In this research, isolation of the coding sequence of plasma membrane Na^+^ /H^+^antiporter (SOS1) and vacuolar membrane Na^+^ /H^+^ exchanger (NHX) in Kochia scparia was performed and, the consequence of salinity stress was studied on the expression profile of this gene. We focused on SOS1 and NHX the critical genes in the SOS pathway and vacuolar membrane for the resistant to salt stress (Fig. 1). The SOS pathway and vacuolar membrane Na^+^/H^+^ exchanger (NHX) are currently the most extensively studied mechanisms in controlling the salt stress response in plants. The SOS and vacuolar membrane Na^+^ /H^+^ exchanger (NHX) pathway is responsible for ion homeostasis and salt tolerance in plants.

### Conserved domains, homology and phylogenetic analyses of SOS1 and NHX in *K.scoparia*

After sequencing the coding SOS1 and NHX genes sequences in *K.scoparia*, Conserved domain specified using of NCBI revealed that putative protein SOS1 belongs to the Sodium/hydrogen exchanger family; These antiporters contain 10-12 trans membrane regions (M) at the amino-terminus and a large cytoplasm region at the carboxyl terminus. The transmembrane regions M3-M12 share the same identity with other members of the which family. The M6 and M7 regions are highly conserved. Thus, this is believed to be the region involved in the transportation of sodium and hydrogen ions. The cytoplasm region has little similarity throughout the family. Conserved domain Analysis for NHX showed that family represents five transmembrane helices. This suggests that the paired regions form a ten helical structure, probably forming the pore, whereas the binds a ligand for export or regulation of the pore. The development of intracellular membrane systems and compartments has led to a considerable increase in the number of ion transporters in eukaryote cells. As a result, plants contain a large number of sequences encoding proteins that share homology to Na/H antiporters which are key transporters in maintaining the pH of actively metabolizing cells. According to highly similar sequences (Mega Blast) search for sequence homology of SOS1 and NHX genes in the NCBI data base on Table3 was provided. Maximum identity up to90% with Suadeamaritimaof NHX gene and for SOS1 gene is 84% by Salicorniabrachiata.

**Table 3:** Analysis sequence alignment using MegaBlast (%identity) related to NHX and SOS1 genes isolated from *K.scoparia.*

| *NHX* |  |  | *SOS1* |  |
| --- | --- | --- | --- | --- |
| No. | Plant species | %Identity | Plant species | Identity % |
| 1 | *Suaeda maritima* | 90% | *Salicornia dolichostachya* | %84 |
| 2 | *Suaeda japonica* | 89% | *Salicornia brachiata* | %84 |
| 3 | *Atriplex gmelini* | 89% | *Suaeda japonica* | %83 |
| 4 | *Suaeda comigulata* | 89% | *Suaeda salsa* | %82 |
| 5 | *Atriplex halimus* | 89% | *Beta vulgaris* | %80 |
| 6 | *Chenopodium glaucum* | 89% | *Spinica oleraceae* | %81 |
| 7 | *Suaeda salsa* | 89% | *Sesuvium portulacastrum* | %79 |
| 8 | *Suaeda pruinosa* | 88% | *Mesembryanthemum crystallinum* | %77 |
| 9 | *Atriplex dimorphostegia* | 88% | *Vitis vinifera* | %78 |
| 10 | *Atriplex patens* | 88% | *Populus trichocarpa* | %76 |
| 11 | *Halostachys scapica* | 88% | *Populus euphratica* | %76 |

According to phylogenic tree; Fig(2) coding sequences SOS1 and NHX genes isolated from *K.scoparia* had the maximum identity with Chenpodiaceae family, for example; SOS1 gene that characterized in *K.scoparia* had maximum homology with Beta vulgaris, Salicorni abrachiata, Spinaceaeoleraceae and suadea salsa,While, show the lowest similarity to Helianthus tuberosus and Sorghum bicolor. As well as a highest similarity NHX gene isolated from *k.scoparia* releated to different species from suaeda genus and the lowest similarity be seen with Brachypodium distachyon.

**Figure 2:**
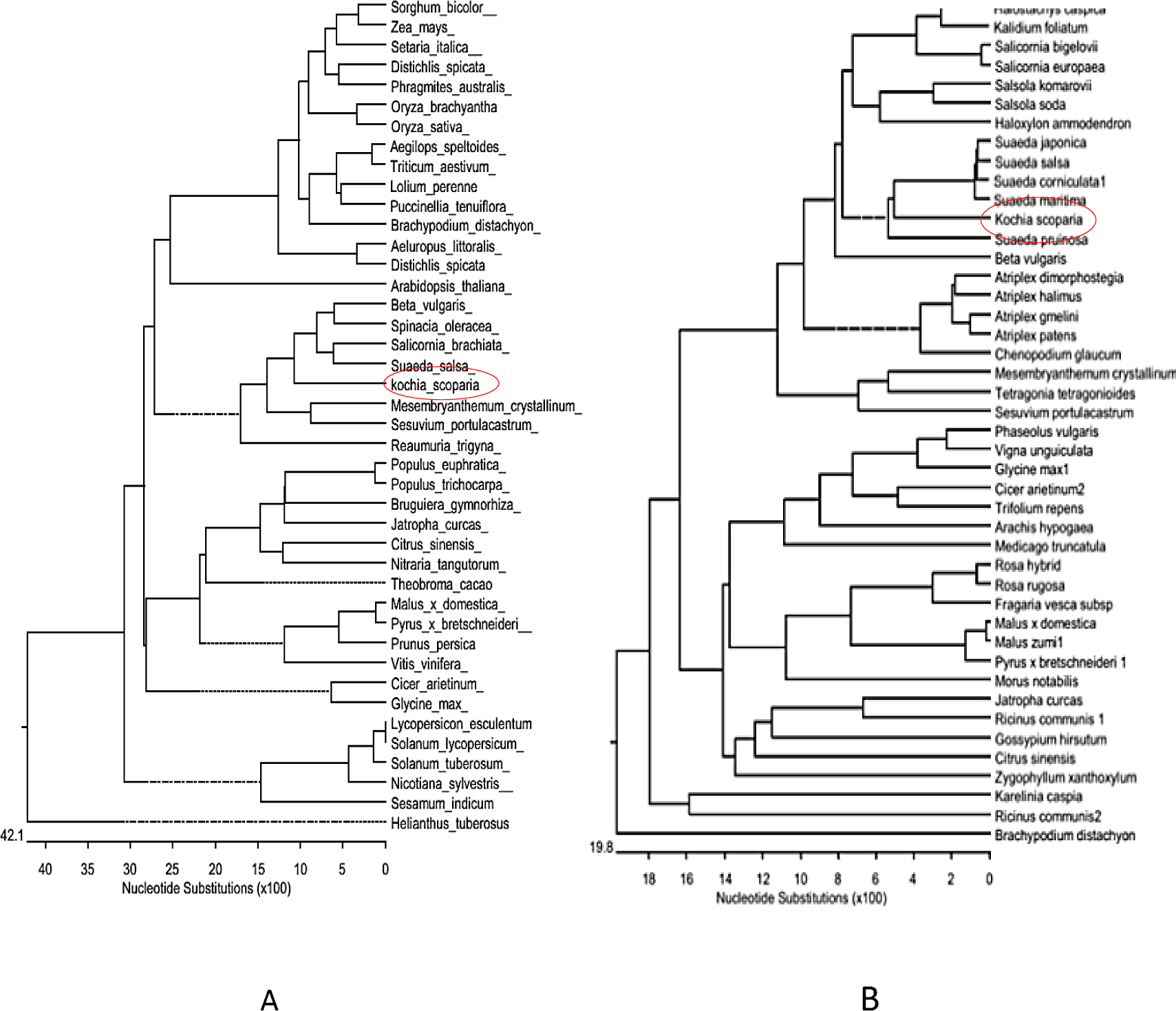
The phylogenic relationship between *K.scoparia* SOS1(A) and NHX(B) with SOS1 and NHX from other plant species. The phylogenic tree was constructed using ClustalW method from the DNASTAR software package. The scale bar indicates substitutions per site. The protein sequences used for construction of the phylogenetic tree are showed on the branch of tree.

### Effects of salinity stress on SOS1 and NHX genes expression profiles

Studies have identified salt tolerance determinants in organisms ranging from cyanobacteria to fungi and from algae to higher plants. Research with halophytic species has provided information on adaptive behavior but information on the molecular level is still Insufficient. Furthermore, information related to salt tolerance of *K.scoparia* at molecular level is insufficient. In this study we tried to be focused on the analysis of isolation, characterization and gene expression pattern of key genes involved in salinity tolerance in halophytes species such as *K.scoparia.* Gene expression profile for SOS1 and NHX were checked in 48hours after treatment with 0, 150, 300 mMNaCl. In this study, we found a basal level of SOS1 and NHX in *K.scoparia* without salt stress, which is regulated with salt treatments. Gene expression Profile for SOS1 and NHX in *K.scoparia* shoot parts showed that salinization was affected SOS1 and NHX levels positively and positive correlation with salinity levels. In other words *K.scoparia* compared to control like most halophytes leaves are progressively increased under all salinity stress. Amounts of mRNA increased for SOS1 gene: 1.5 and 2.5 and NHX gene: 1 and 2 times higher than the control (0mM) in 150 and 300mM stressed plants after 48 hours of exposure respectively (Fig.3). While amounts of mRNA increased for SOS1 gene and NHX in root plant but less than the increase in leaves, 1 and 2 times for SOS1, 0.5 and 1 fold higher than the control in 150 and 300mM treated plants (Fig.3).

**Figure 3:**
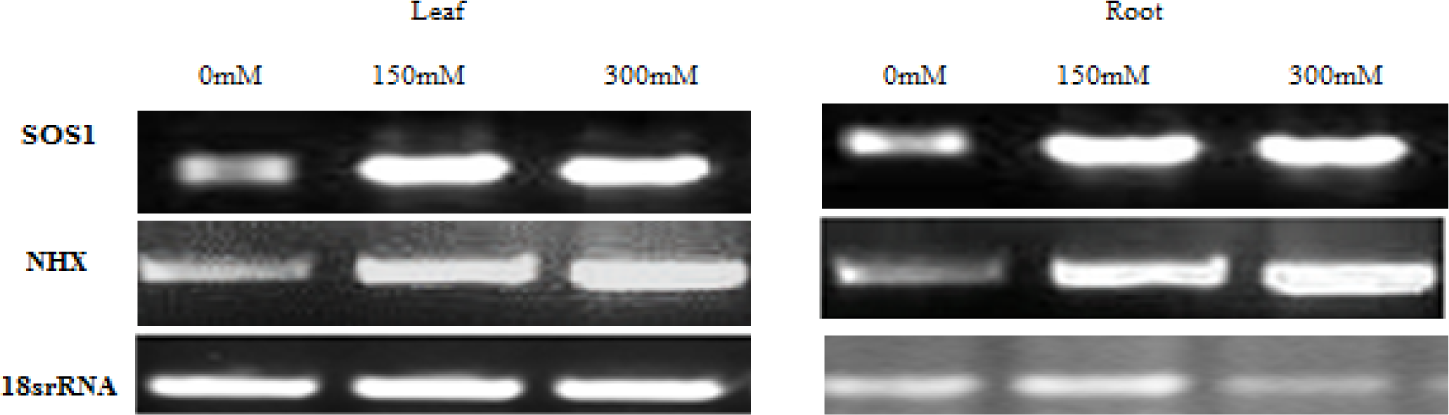
Semi quantitative RT-PCR analysis showing differential gene expression in leaf tissues and roots of 12-day-old seedlings for SOS1 and NHX gene of *K.scoparia.* The expression of each gene was compared relative to its expression in control gene (18s rRNA). Samplings were carried out at 24 hours after treatments with 0, 150, 300 mM salt stress.

### Prediction of SOS1 antiporter using Bioinformatic tools

For prediction binding of Cyclic nucleotide to cyclic nucleotide-binding domain to SOS1 protein isolated from *K.scoparia:* First, the tertiary structure of the desired protein domain was predicted (Automated Mode, The pipeline will automatically identify suitable templates based on Blast (Altschul, Gish et al. 1990) and HHblits (Remmert, Biegert et al. 2012). Cyclic nucleotide-binding domain in these proteins has a For prediction binding of Cyclic nucleotide to cyclic nucleotide-binding domain to SOS1 protein isolated from *K.scoparia*: First, the tertiary structure of the desired protein domain was predicted (Automated Mode, The pipeline will automatically identify suitable templates based on Blast and HHblits. Cyclic nucleotide-binding domain in these proteins has a common structure with 120 amino acids. The domain consists of three alpha helixes and eight turn structures that form a porelike structure. Three protected glycine amino acids seem important to maintain the barrel-shaped structure. In the anticipation of the domain in the *K.scoparia*, structures such as antiparallel beta sheets and alpha helix associated with small screws. Using molecular docking (using by SwissDock, a protein-small molecule docking web service based on EADock DSS (Kumar, Kumar et al. 2013)) of the desired fragments separated from Kochia plant as shown in Figure (4). Figure 4, the pore-like structure created in the first part of cyclic nucleotide binding site, provides the most likely connection for cyclic nucleotide cNMP.

**Figure 4:**
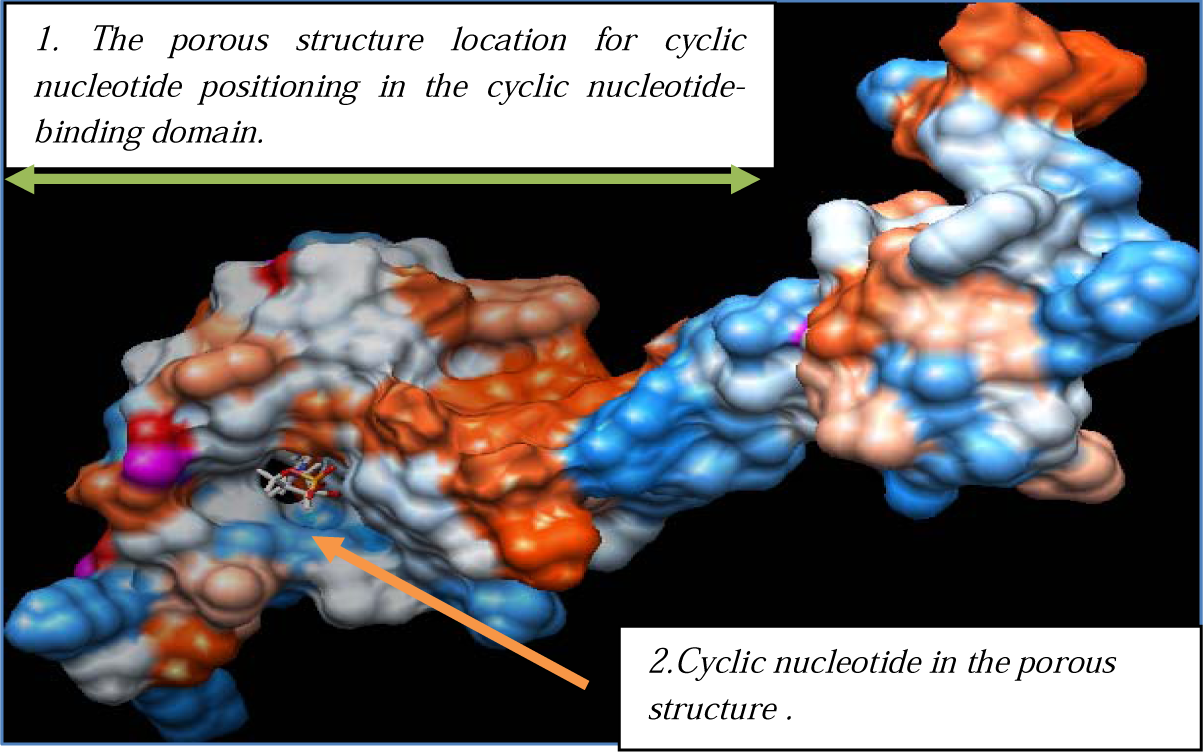
Molecular docking was performed to locate the meeting point of cyclic nucleotide binding to SOS1 protein separated from *K.scoparia* using the online service of Swiss Dock and UCSF-Chimera software. The placed mark on Fig is the pore-like structure available in the area for suitable connection between cyclic nucleotide and the desired location in SOS1 protein isolated from *K.scoparia.*

## Discussion

Studies have identified salt tolerance determinants in organisms ranging from cyanobacteria to fungi and from algae to higher plants. In plant cell maintain a high K^+^/Na^+^ in the cytoplasm, under normal conditions. Under salt stress conditions plants have several strategies and adaptive mechanisms for tolerant to these conditions. In these mechanisms, to be launched sensing, signal transduction, gene expression and metabolic pathways. Evidence may be these tolerance progarms slow and steady adapatation in the sensitives plants. Therefore, understanding the components of these mechanisms in halophytes can do contribute substantially to improving retrofitting sensitive plants.We focused on isolation, characterization and gene expression pattern analysis of main genes involved in salinity tolerance in halophytes from *K.scoparia.* In the present study, the relatively high basal expression level of V-NHX indicated the important physiological function of NHX in *K.scoparia*, even in the absence of stress NHX levels positively have correlation with salinity levels. In other words, compared to control *K.scoparia* like most halophytes leaves progressively increases under salinity stress. Amounts of mRNA increased for NHX gene: 1 and 2 times higher than the control (0mM) in 150 and 300mM stressed plants after 48 h of exposure respectively (Fig. 4), The higher NHX1 expression in the leaves was a prompt response to NaCl treatment which could have helped decrease the Na^+^ content in the cytoplasm and maintain water concentrations (Khedr, Serag et al. 2011).Previously transgenic studies have shown that the over expression of the NHX1 gene significantly enhanced plant salt tolerance abilities, in transgenic Arabidopsis over expressing AtNHX1, higher activities of the vacuolar Na^+^/H^+^ antiporter were observed and enabling it to grow in the presence of 200 mMNaCl (Apse, Aharon et al. 1999). The over expression of the cotton Na^+^/H^+^ antiporter gene GhNHX1 in tobacco improved salt tolerance in comparison with wild-type plants (Wu, Yang et al. 2004). Na^+^/H^+^ antiporter is an important membrane protein responsible for pumping Na^+^ into the vacuole to reduce Na^+^ toxicity and alleviate the adverse effects of salt stress (Wu, Yang et al. 2004).The expression of the K.s SOS1 gene in L. fusca was regulated by Na ^+^ and to characterize the engagement of SOS1 in kochia response to saline conditions, results showed that a basal level of *K.scoparia* SOS1 transcripts in plants without salt stress, which up-regulated significantly with salt treatments. amounts of mRNA increased for SOS1 gene and NHX in roots plant but less than the increase in leaves, once and towice times for SOS1, 0.5 and 1 time higher than the control in 150 and 300mM treated plants. These results are in agreement with the Oh et al. 2009, report about the increase of the SOS1 expression level in response to salt treatments in A. thalianaand T. halophila. It has been reported that A. thaliana AtSOS1is expressed at low basal levels but is up-regulated significantly by salt stress in both roots and shoots. Moreover, it regulates other genes in response to salt stress (Shi et al. 2000, Gong et al. 2001). Based on the conducted docking; there is a possibility of hydrogen and hydrophobic connection in the porous structure. The amino acid phenylalanine, lysine, threonine, glycine and arginine have the most connection and positioning in the porous structure according to their charge, polar, non-polar and structure features. Meanwhile, the conserved glycine amino acids, which are involved in the formation of pore-like structure, are also effective in hydrophobic connections,With further investigation on this domain, these factors affecting it can be determined. In the other plants antiporter activity, long cytosolic C-Terminal tail of SOS1 in thought to be involved in the sensing of Na^+^ (She et al., 2000). Furthermore SOS1 has been demonstrated to be a target of SOS pathway, releationships between SOS1 and SOS2/SOS3 can be a way of regulating the activity of SOS1. Ceratin domains of SOS1 reacted with SOS2/SOS3 that charectrization of these domains can be helped to the use of this protein in process will creat resistance plants. To provide factors will be affected the SOS1 activity and to determine suitable methods to activate these proteins in Glycophyt plant. On the other NHX exchanger acts as a mediator of K^+^ taransport between cytosol and vacoule, SOS2 also activates the vacuolar-ATP as e and vacuolar Na^+^/K^+^ antiporter NHX exchangers, which compartmentalizations Na^+^/K^+^ into vacuoles. K^+^ compartmentalization in the vacuole coulad result in a cytosolic K^+^ deficiency(Zhang et al., 2014). So determine how to communicate this antiporters can be controlled out ways of increasing salt tolerance to be properly. Sequencing of SOS2 and prediction of interact with NHX can be used for controling of SOS pathway.

## Conclusion

In the my study SOS1 and NHX genes sequenced and determind Proteins characteristics with insilico tools. Charecterization of other genes involved these pathways and signaling pathway and investigation invitro of proteins are apromising area of research that may lead to improvments in the biomass production of crop with external applications materials and genetic manipulation.

